# Proteomic Characterization of Prostate Cancer Bone Metastasis Subtypes Defines Progressive Remodeling States of the Bone-Tumor Microenvironment

**DOI:** 10.64898/2026.10.08.757504

**Authors:** Dennis Xie, Bita Mojtahedzadeh, Keanu Chan, Charmaine Chui, Hans Adomat, Naomi Jung, Doris Liang, Colm Morrissey, Eva Corey, Lawrence D. True, Rizhi Wang, Felipe Eltit, Michael E. Cox

**Affiliations:** Vancouver Prostate Centre, Vancouver, Canada; Department of Urologic Sciences, University of British Columbia, Canada; School of Biomedical Engineering, University of British Columbia, Canada; Centre for Aging SMART, Vancouver, Canada; Department of Urology, University of Washington, USA; Department of Laboratory Medicine and Pathology, University of Washington, USA; Department of Materials Engineering, University of British Columbia, Canada

## Abstract

Prostate cancer (PC) bone metastases (PCBM) are a debilitating disease morbidity that primarily affects the axial skeleton. PCBM are predominantly osteosclerotic with mixed osteosclerotic/osteolytic features, including irregular bone deposition, loss of collagen alignment and increased porosity that mimic features of woven bone formed during development and repair. We hypothesize that progression of PCBM is driven by interactions between tumor cells and the bone microenvironment and is associated with changes in the stromal compartment. We performed tandem mass tag mass spectrometry proteomics on lumbar vertebral specimens from 32 cadaveric PCBM specimens and 3 age-matched controls. Unsupervised clustering segregated specimens into three clusters associated with benign, mixed, and osteosclerotic lesion states, characterized by distinct immune-, tumor-, and osteogenic-associated protein signatures. Histological analysis showed tumor infiltration was accompanied by marrow fibrosis and increased collagen III deposition. We identified a marker signature associated with osteosclerotic remodeling and mapped this signature to independent metastatic PC transcriptomic datasets. The proteomics-derived markers reproduced similar molecular groupings at RNA level, and robustly segregated bone PCBMs from soft tissue metastases PC in RNA data. Together, these findings define molecularly distinct states of tumor-driven changes in PCBM that extend beyond histological classification and may inform biomarker development and therapeutic stratification.

## Introduction

Prostate cancer (PC) is the second most commonly diagnosed malignancy and the fifth leading cause of cancer-related death among men worldwide, accounting for more than 1.46 million new diagnoses and approximately 396,000 deaths in 2022 (1). Despite the success of modern therapies, 14% of patients progress to incurable metastatic castration-resistant PC and approximately 90% of these patients will develop bone metastases (2, 3).

PC bone metastases (PCBM) are a devastating condition that affects the hematopoietic skeleton of spine, ribs, and hips (4, 5). PCBM are associated with a variety of bone alterations from increased bone density (osteosclerotic) to reduced bone density (osteolytic). These lesions increase fracture risk, generate excruciating pain, and dramatically affect patients’ quality-of-life (2). Structurally, osteosclerotic PCBM lesions present with irregular bone distribution, loss of collagen alignment, and increased porosity mimicking features of woven bone that form during development and repair (6). Patients who fail primary therapy, and patients diagnosed with advanced disease face prolonged treatment with androgen receptor (AR) pathway inhibitors (7–9). These agents are primarily responsible for improved survival, but as men live longer with the disease, the clinical management of time-associated complications, especially the cumulative burden of bone metastatic disease, has become increasingly important (7, 10).

Mundy and Guise’s “Vicious Cycle” hypothesis postulates that PC cell-presence in the bone environment leads to dysregulated bone homeostasis and induction of pathologic bone architecture changes (11, 12). Accelerated bone turnover releases growth and survival factors supporting adaptive changes in PC that affect disease pathogenesis. By analogy to Dvorak’s “Tumors: wounds that do not heal” hypothesis, we reported that PCBM bone undergoes a similar process, wherein reparative (woven) bone is produced, but never transitions to mature bone (13). This woven bone has higher porosity, irregular collagen organization, and prominent collagen III (COL III) content consistent with a compromised bone structure. While collagens are the most abundant bone proteins that form the mineralized structural scaffold of the skeleton, many other proteins and proteoglycans are critical for bone homeostasis by modulating mineral deposition, directing collagen distribution, and allowing cell adhesion and survival (14–16). Thus, changes to the extracellular matrix (ECM) composition can alter bone structural and mechanical properties. Despite these characteristic pathological changes, the molecular composition of PCBM bone and stromal compartments remains poorly understood as previous proteomic and transcriptomic studies have largely focused on tumor cell features, identifying distinct molecular subtypes associated with androgen receptor signaling, proliferation, DNA-repair activity, and stromal–epithelial interactions (17, 18).

In this study, we characterized three distinct proteomic ECM phenotypes in PCBM. These phenotypes reflected progressive remodeling of the metastatic niche, including loss of native immune and hematopoietic architecture in the bone marrow, an expansion in tumor associated signatures, and increased osteogenic activity. Contrasting these results through IHC further revealed marked ECM remodeling, including excessive collagen I (COL I) and COL III deposition in the marrow space and the presence of bone remodelling markers bone sialoprotein (IBSP) and osteopontin (SPP1) in both sclerotic bone and tumor cells. We observed similar phenotypic patterns, and confirmed increased osteogenic and ECM remodeling signatures, in publicly available RNA sequencing (RNA-seq) datasets derived from metastatic tumor biopsies from metastatic castration resistant PC (mCRPC) patients. Our analysis is the first to characterize changes in the bone microenvironment protein composition. By identifying representative features of a tumor-driven bone remodeling continuum in PCBM, we establish protein signatures that may inform biomarker development and targeted therapeutic strategies.

## Results

### Proteomic profiling defines subgroups of PCBM with distinct histological features

Bone is predominantly composed of a mineralized ECM. To characterize any changes in the ECM protein composition of PCBM, we performed tandem mass tag (TMT)-based mass spectrometry (MS) proteomics profiling on PC cadaveric vertebrae and age-matched controls. We performed micro-computed tomography (microCT) scans on a total of 67 PCBM specimens, and selected the 15 most osteolytic and the 17 most osteosclerotic samples for proteomic analysis, and compared the results to analysis of 3 age-matched control bone specimens (6). It is important to note that microCT classification reflects the architecture of the entire bone core, but it may not accurately represent localized tissue heterogeneity in regions subjected to proteomic analysis. We, therefore, quantified BV/TV of histological sections sequential to the analyzed regions and to confirm PCBM microarchitecture subtype classification and the presence of tumor.

Based on our TMT-MS approach, we detected 888 proteins across all samples, of which 412 were detected in more than 50% of the samples. From the 412 proteins we observed three distinct clusters from the principal component analysis (PCA) (**Figure 1A**). Sample clusters 1 and 2 exhibited low mean BV/TV (0.109 ± 0.035 and 0.130 ± 0.037; mean ± SD, respectively), whereas cluster 3 displayed high BV/TV (0.608 ± 0.148) (**Figure 1B, Supplemental Table 1**). Cluster 1 (n= 8) consisted exclusively of age-matched control specimens from individuals without prostate cancer and histologically benign specimens from patients with metastatic prostate cancer, in which no tumor cells were identified in the analyzed bone sections. Sirius Red staining of samples in this cluster demonstrated preserved trabecular organization with intact marrow spaces and no evidence of aberrant bone deposition (**Figure 1, C and F and Supplemental Figure 2 A**). Cluster 2 (n= 11) contained a mixture of benign and tumor-containing specimens with BV/TV values ranging from 0.082 to 0.204. Although BV/TV remained low in this cluster, Sirius Red staining revealed abnormal ECM remodeling in the tumor-containing specimens, including areas of deposition of woven bone and encrustation along trabecular surfaces with areas of osteolytic activity signified reduced trabecular thickness and increased void space **(Figure 1, D and G and Supplemental Figure 2 B**). Cluster 3 (n= 16) contained exclusively tumor-containing samples and is distinctly characterized by elevated BV/TV (0.608 ± 0.148). Sirius Red staining showed these specimens were predominantly osteosclerotic and included both lesions with residual trabecular structure and lesions lacking discernible trabeculae, consistent with previously described osteosclerotic subtypes (6) (**Figure 1, E and H and Supplemental Figure 2 C)**.

**Figure 1.**
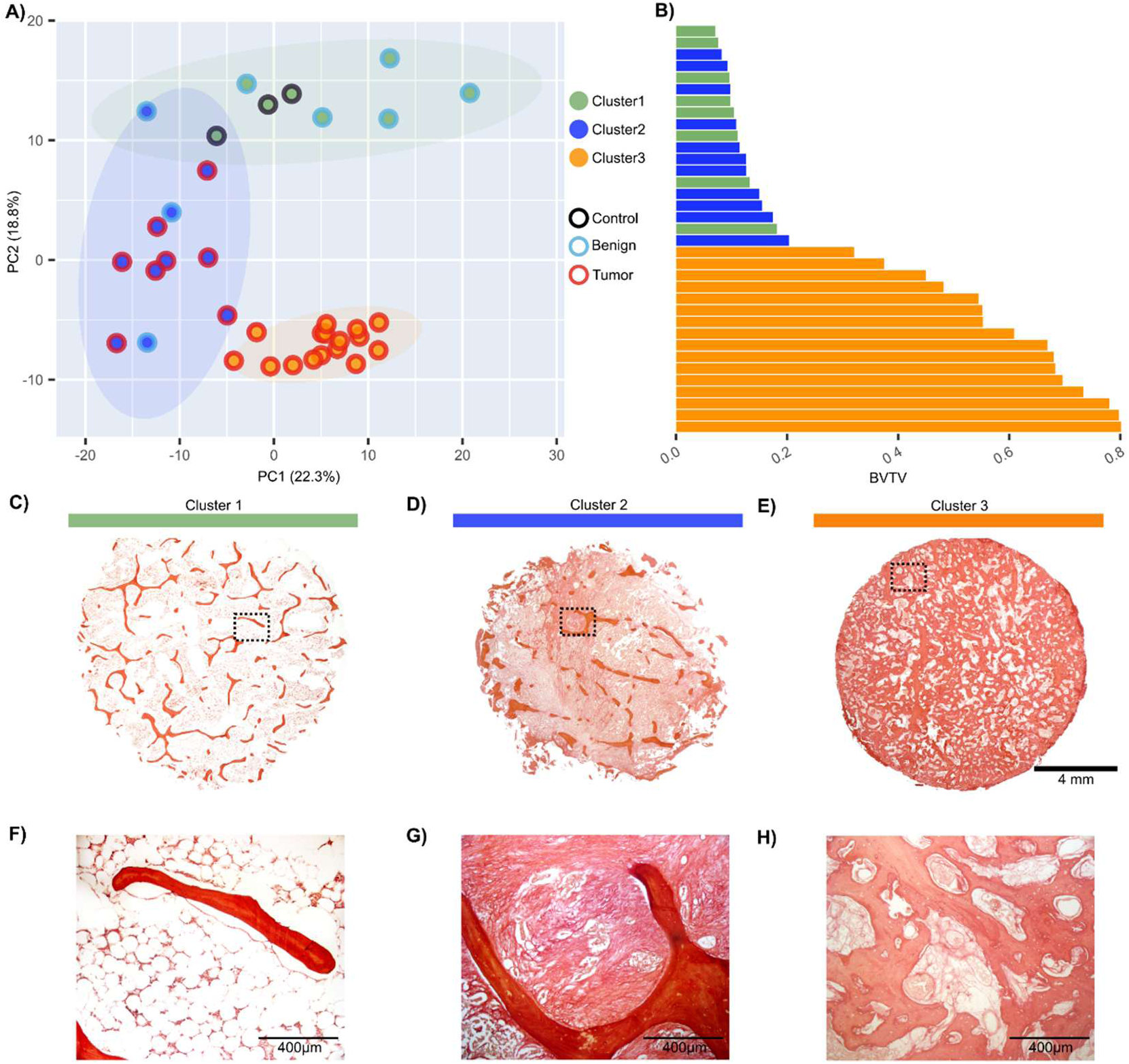
Proteomic profiling defines subgroups of PCBM by lesion types. (A) PCA of the proteomic profiles. Each dot represents an individual sample; dot and ellipse colors indicate K-means clustering results. Ellipses denote 95% confidence intervals. Controls are shown in dark green. (B) Waterfall plot BV/TV measurements for each sample, calculated from sequential Sirius Red (SR) sections. (C–E) Representative SR-stained sections from Cluster 1 (C), Cluster 2 (D), and Cluster 3 (E). Black boxes indicate regions shown at higher magnification. (F–H) Corresponding 10× magnification images of the boxed regions in C–E

To assess differences in bone marrow architecture and tumor involvement, we examined H&E-stained sections from samples within each cluster **(Figure 2)**. Samples in cluster 1 retained intact hematopoietic marrow architecture in both control specimens (**Figure 2A**) and histologically benign PCBM specimens (**Figure 2B**). While samples in cluster 2 preserved areas of hematopoietic architecture, clear tumor infiltration was observed in the marrow space in 8 of the 11 samples (**Figure 2C**). These tumor-containing specimens also exhibited substantial fibrosis. In cluster 3, tumor infiltration was present in all 13 specimens and was associated with extensive osteosclerotic bone formation and a complete loss of normal hematopoietic bone marrow (**Figure 2D**).

**Figure 2.**
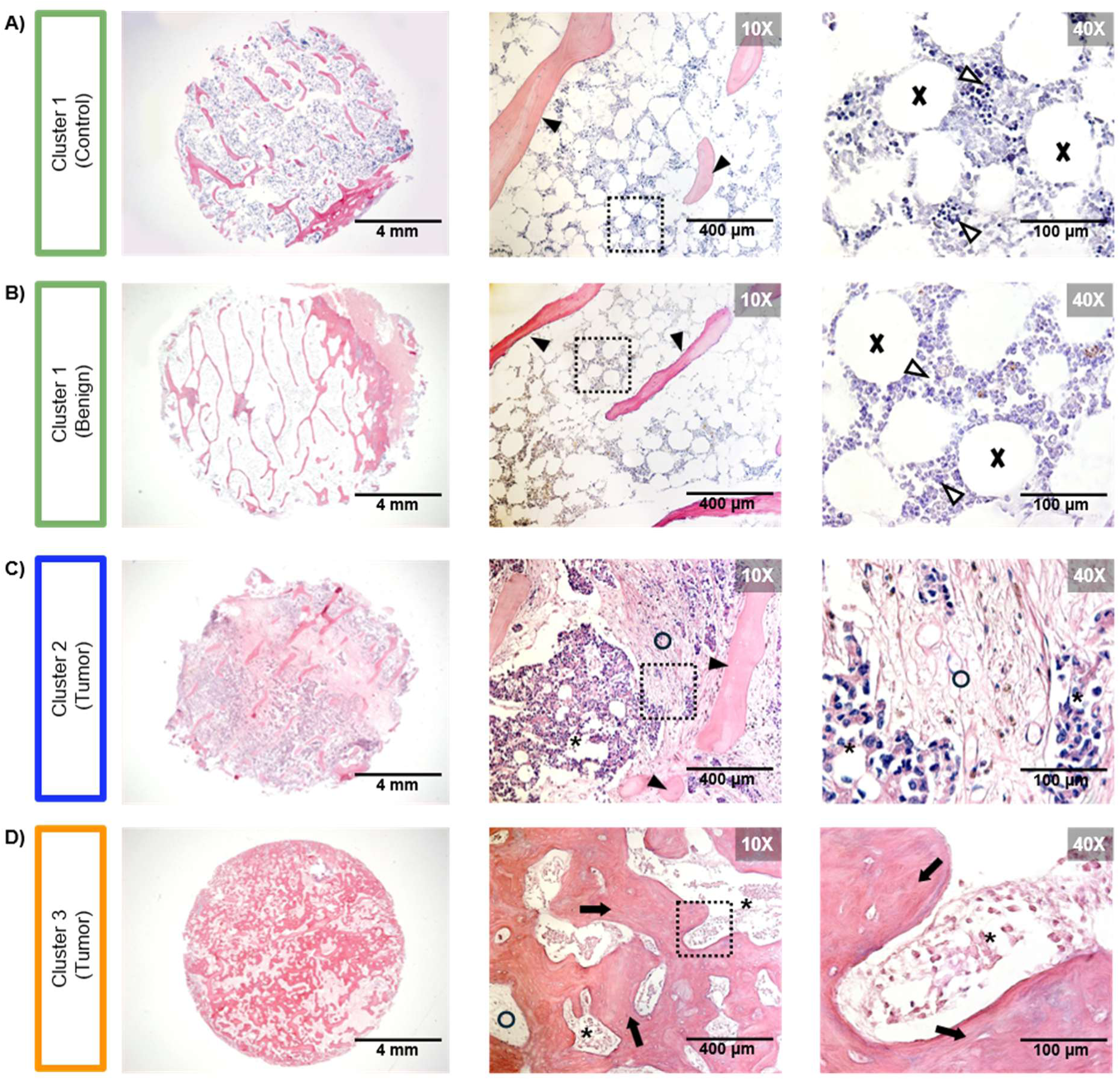
Altered marrow architecture in PCBM. (A-D) H&E staining of representative specimens from cluster 1-3. Left panels show low-magnification H&E overviews of representative specimens. Middle panels show corresponding 10× magnification views, and right panels show 40× magnification views of the regions indicated by black boxes. (A) Cluster 1 control bone marrow shows preserved hematopoietic elements (open arrowheads), trabecular bone (arrowheads), and adipose tissue (X). (B) Cluster 1 histologically benign bone marrow. (C) Cluster 2 mixed lesion with tumor cells (asterisks), fibrotic stroma (open circles), and residual trabecular bone (arrowheads). (D) Cluster 3 osteosclerotic lesion with tumor cells (asterisks) and osteosclerotic bone matrix (arrows).

### Proteomics analysis reveals functional differences between clusters

To identify underlying pathways and markers contributing to cluster separation, we performed pairwise differential expression protein (DEP) analysis between the three clusters using a threshold of ≥ 1.5-fold change and adjusted p ≤ 0.05. Cluster 1 contained 57 uniquely upregulated proteins relative to clusters 2 and 3. Only 9 of the differentially expressed proteins overlapped with cluster 3, and none overlapped with cluster 2 (**Figure 3A)**. Cluster 2 contained 34 uniquely differentially expressed proteins, and cluster 3 contained 39. These DEP analyses indicate that the bone microenvironment is substantially altered across the disease spectrum and represented by these clusters. Notably, cluster 2 and 3 shared 50 differentially expressed proteins, suggesting a shared continuum of bone metastatic niche changes across these two clusters. From these comparisons, a total of 58 differentially expressed proteins were identified as cluster-specific marker proteins, with each marker uniquely upregulated in one of the three clusters.

**Figure 3.**
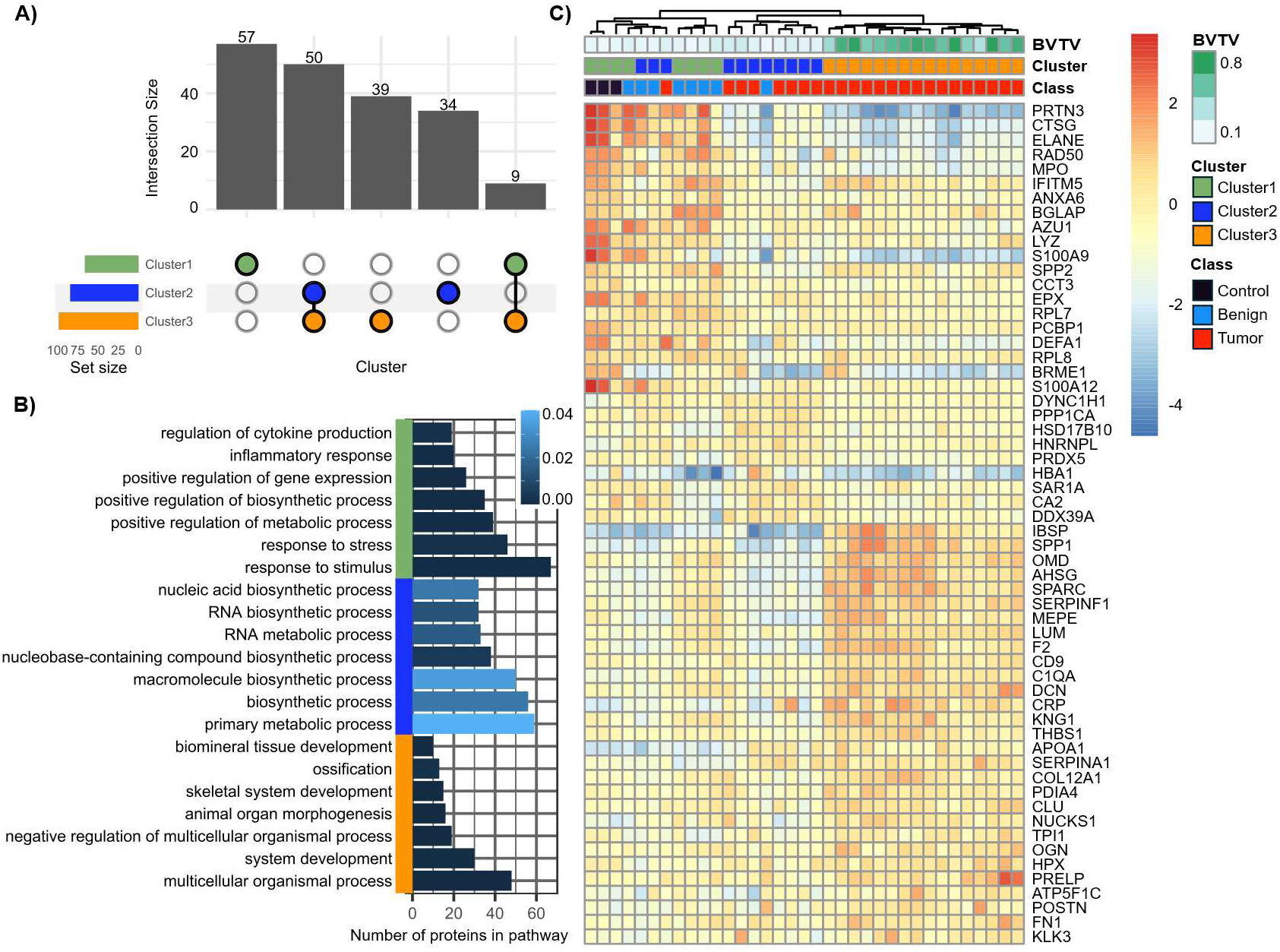
Differential expression analysis showed functional differences between each cluster. (A) UpSet plot showing intersections between upregulated proteins from each cluster comparison. Horizontal bars indicate the total number of cluster-associated proteins identified for each cluster. Vertical bars indicate the number of proteins unique to or shared between clusters, with connected filled circles below each bar denoting the corresponding cluster intersection. (B) Functional enrichment analysis of protein groups identified by hierarchical clustering. Horizontal bar color indicates significance. (C) Relative protein expression of targets uniquely upregulated in each cluster. Color bars represent BV/TV, sample cluster and samples type.

To determine the functional implications of these proteomic changes, we performed overrepresentation pathway analysis with gProfiler using DEP profiles from each cluster. Cluster 1 was enriched for immune-related pathways, including innate immune signaling and inflammatory processes (**Figure 3B**), suggesting preservation of bone marrow immune niches in control and benign bone. Cluster 2 was enriched for cell metabolism and proliferation pathways, consistent with a tumor-establishment state characterized by active cellular remodeling, metabolic activity, and proliferation. Cluster 3 showed strong enrichment for extracellular matrix organization, skeletal system development, and bone turnover pathways, indicating activation of osteogenic and matrix-remodeling programs under the influence of PC cells. Although histologically benign samples generally clustered with cancer-free controls, their proteomic profiles were not uniformly consistent with those of the control group. Notably, 3 benign samples were associated with Cluster 2, and showed slightly elevated PSA levels despite the absence of detectable tumor invasion by routine histology or immunohistochemistry (**Figure 3C**). Because the proteomic profile of these samples matched tumor-containing samples in cluster 2 but did not match the control samples in cluster 1, this discrepancy may indicate that proteomic profiling can detect early or spatially restricted changes that occur below the threshold of conventional histopathological detection. We profiled an additional 19 samples using the same TMT-MS workflow and performed PCA on the combined sample group using the 44 proteins overlapped with the 58 marker proteins. The original specimens retained the same clustering pattern as in the initial analysis, the additional samples followed similar trends when cross validating histological characteristics and PCA clustering, with one exception where a control sample was grouped into cluster 2 (**Supplemental Figure 3**).

To identify specific proteins upregulated in each cluster we performed pairwise cluster comparisons. The full list of quantified proteins, abundances, and fold changes for each comparison is provided in the Supplemental Data. In Cluster 1, highly upregulated proteins included PRTN3 and CTSG, both neutrophil-associated serine proteases, along with MPO and ELANE, both markers of granulocyte maturation, together indicating preserved hematopoietic differentiation and immune function (**Figure 4, A and B**). In cluster 2, upregulated proteins include ARF3 a small GTP-binding protein involved in vesicle binding which have been implicated to play a role in PC metastasis and CMPK1 an important protein in nucleotide phosphorylation (19) (**Figure 4A**). In cluster 3, CRP, APOA1, APOA2 and tumor-associated protein TPD52 are upregulated compared to cluster 1 (**Figure 4B**). In parallel, Cluster 3 also showed enrichment of bone matrix proteins IBSP and SPP1, which are non-collagenous proteins involved in mineralization, osteoblast/osteoclast interactions, and tumor–bone adhesion, while SPARC is a matricellular protein involved in collagen organization and extracellular matrix remodeling (20) (**Figure 4, B and C**). The Cluster 3 upregulated proteins reflect both a diseased-state and osteogenic bone remodelling phenotype.

**Figure 4.**
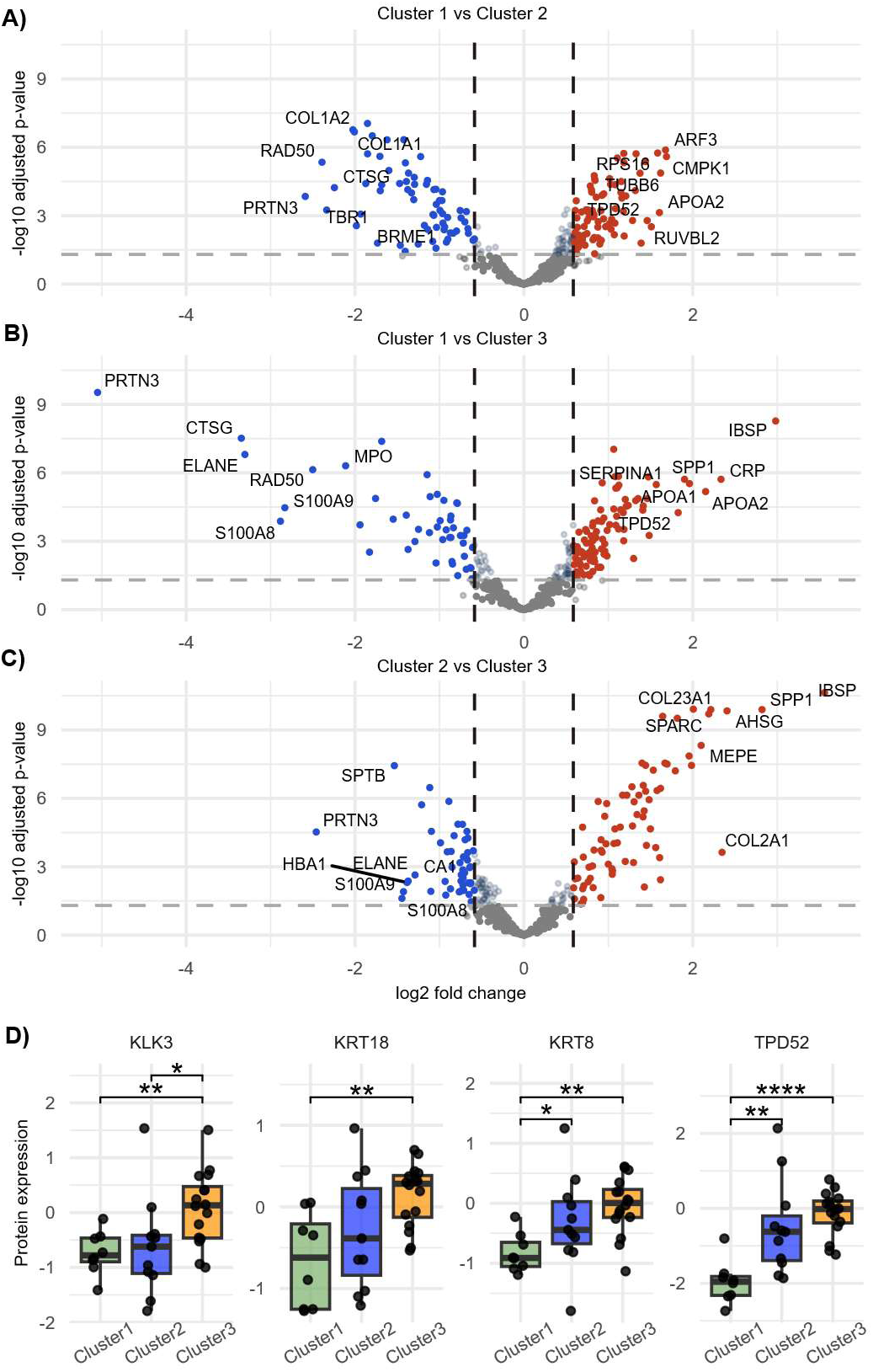
**(A-C)** Volcano plot showing top 7 differentially expressed proteins for each pairwise cluster comparison. (A) Cluster 1 vs Cluster 2, (B) Cluster 1 vs Cluster 3, (C) Cluster 2 vs Cluster 3. (D) Boxplot showing average expression level of PC-associated proteins across clusters. \**p* < .05, \*\**p* < .01, and \*\*\*\**p* < .0001.

To determine whether differences in PC content contributed to the proteomic separation of the clusters, we independently examined the expression of the four PC-associated proteins consistently detected across all samples (**Figure 4D**). KLK3 (prostate-specific antigen, PSA) expression was elevated in cluster 2 relative to cluster 1 and was highest in cluster 3, while TPD52 was also significantly increased in clusters 2 and 3. Both KRT8 and 18 are significantly unregulated in cluster 3 compared to cluster 1. Combined, these results revealed an underlying shift in the bone microenvironment that accompanied by the emergence of PC cells.

### Cluster-associated expression reveals osteogenic and fibrotic remodeling in PCBM

From the differential expression analysis described above, IBSP and SPP1 showed the largest fold increases in Cluster 3. Both proteins are key regulators of bone remodeling and mineralized matrix formation (20). We performed IHC to validate their spatial distribution within osteosclerotic PCBM lesions. In control bone, staining of IBSP, a marker of osteogenic activity and bone remodeling, was largely restricted to osteocytes in the trabecular bone (**Supplemental Figure 4**) (14). In contrast, PCBM specimens demonstrated stronger IBSP expression in regions of woven sclerotic bone compared to mature trabecular bone (**Figure 5, A-D**). Intense IBSP staining was observed in osteoblasts lining the surfaces of woven bone and in osteocytes embedded within the newly formed matrix, whereas osteoblasts associated with organized trabeculae exhibited comparatively weaker staining (**Figure 5, A-D**). Additionally, strong staining can be observed in the tumor cells within the bone microenvironment, consistent with previously described osteomimetic features of PC (21). SPP1 displayed a similar spatial distribution, with strong expression in the tumor and osteoblasts and osteocytes associated with woven bone (**Figure 5, E-H**). However, unlike IBSP, SPP1 staining within the woven bone matrix itself was less prominent, suggesting that although both proteins are associated with pathological bone formation, they occupy different extracellular niches during osteogenic remodeling.

**Figure 5.**
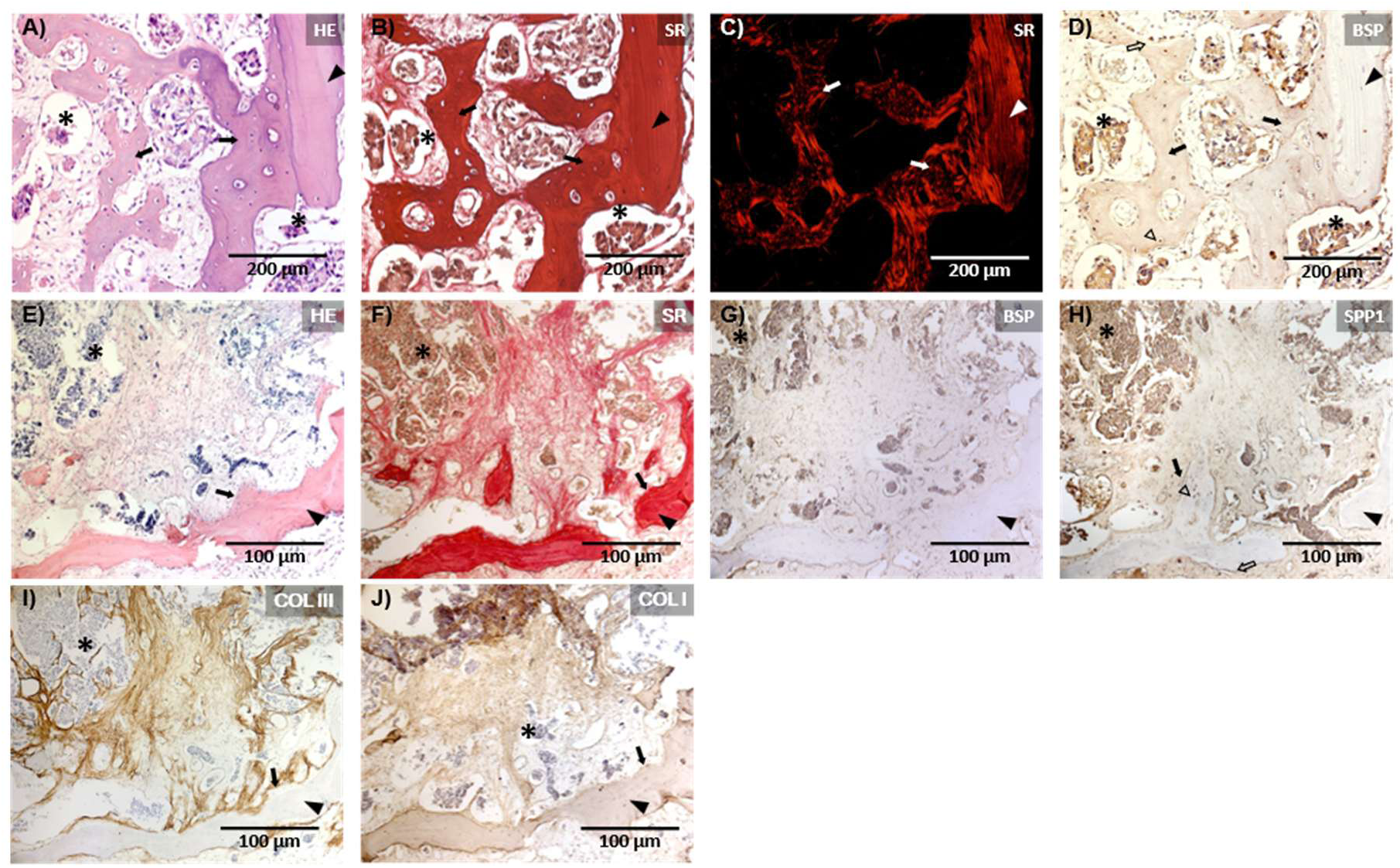
Dysregulated expression of ECM proteins in PCBM. (A) H&E staining of PCBM showing residual trabeculae bone (arrowheads), osteosclerotic bone (arrows), and tumor cells (asterisks). (B) Sirius red (SR). (C) Polarized light imaging of B. (D) IBSP in osteosclerotic bone (arrows), osteocytes (open arrowhead), osteoblasts (open arrow) and tumor cells (asterisks). (E) H&E staining of a region containing newly formed bone adjacent to fibrotic marrow. (F) Sirius Red staining corresponding to E. (G) IBSP staining. (H) SPP1 staining. (I) COL III staining. (J) COL I staining.

Bone marrow fibrosis and osteosclerosis are closely linked in bone marrow disease like primary myelofibrosis (22). To evaluate fibrotic extracellular matrix remodeling, we assessed COL III deposition. COL III was deposited as a matrix admixed with COL I. This admixed matrix was observed surrounding tumor nests and accumulating at the interface between tumor cells and woven bone (**Figure 5, E-J and Supplemental Figure 5, A-C**). In addition, COL III formed a dense fibrillar meshwork throughout affected marrow spaces (**Figure 5, I and Supplemental Figure 5, D-I**). While COL I fibers can also be observed in these spaces, by comparison, COL III localized predominantly to the unmineralized fibrotic matrix surrounding newly formed bone, whereas mineralized bone matrix itself was COL III-negative and COL I positive (**Figure 5, I and J**). Based on these observations, we conclude that PCBM specimens with a prominent cancer cellular component exhibited extensive fibrosis characterized by active deposition of immature woven bone enriched for osteogenic markers alongside extensive fibrotic remodeling consists of COL I and COL III fibers, consistent with activation of tissue repair and pathological bone remodeling pathways.

### PCBM marker panel reveals transcriptional PCBM subtypes with varying immune and osteogenic signatures

To determine whether this protein-based segregation of PCBM phenotypes was also reflected by transcriptional profiles of metastatic lesions, we analyzed PCBM samples from the merged set of publicly available RNA-seq datasets: the Stand Up 2 Cancer/Prostate Cancer Foundation (SU2C/PCF) East Coast Dream Team (ECDT), West Coast Dream Team (WCDT), and Prostate Cancer Medically Optimized Genome-Enhanced Therapy (PROMOTE) cohorts (**Figure 6A**). We used the 58 proteomics-derived cluster markers as a feature set and performed independent unsupervised clustering of the transcriptomic data. We then compared the expression patterns and biological characteristics of the resulting transcriptomic groups with the proteomic clusters to determine whether similar molecular states were recapitulated across data modalities. Unsupervised clustering divided the PCBM RNA data into 3 groups, RNA cluster A-C (**Figure 6B**). PCA was further used to demonstrate clear separation of the three transcriptional groups, consistent with their unsupervised cluster assignments (**Figure 6C**). Gene set enrichment analysis (GSEA) analysis of the three transcriptional groups demonstrated patterns resembling the proteomic clusters. One cluster was enriched for immune-related pathways, one for cell metabolism and proliferation-associated pathways, and one was enriched for extracellular matrix organization and bone remodeling pathways (**Figure 6D**). These findings suggest that the proteomic phenotypes identified in our cohort are recapitulated at the transcriptomic level across independent bone metastasis datasets. To determine whether this molecular classification was specific to bone metastases, we applied the same marker panel to transcriptomic data from non-osseous metastatic sites within the integrated dataset (**Supplemental Figure 6, C and D**). In contrast to bone metastases, samples from other metastatic sites did not segregate into the same three molecular groups, suggesting that these phenotypes are specific to protein expression changes in the PCBM microenvironment.

**Figure 6.**
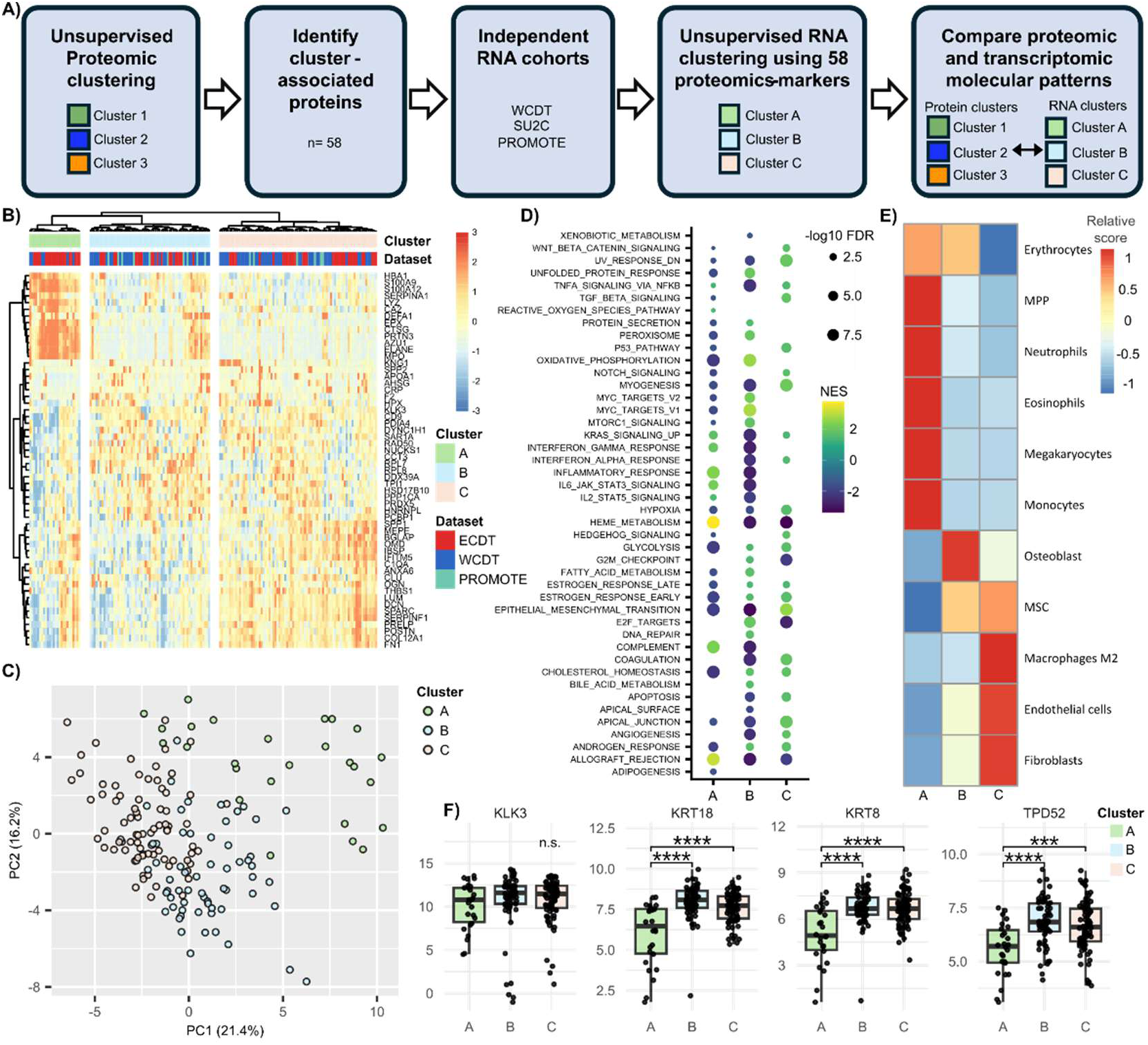
Transcriptional PCBM subtypes with varying immune and osteogenic signatures. (A) Workflow illustrating the application of proteomics-derived markers to independent transcriptomic datasets. (B) Unsupervised hierarchical clustering heatmap of proteomics-derived marker genes in BM samples from the merged RNA-seq dataset. Samples were separated into three transcriptional clusters by cutting the hierarchical clustering dendrogram generated using correlation distance. Column annotations indicate RNA cluster and dataset. (C) PCA plot showing distribution of samples by the top 1500 variable genes, color indicates cluster. (D) GSEA analysis of hallmark pathways in each of the RNA transcriptional clusters. Color of the circle indicates normalized enrichment score (NES) and circle size represent -log10 false discovery rate (FDR). (E) Cell type enrichment score across clusters. (F) Expression of prostate cancer markers across clusters. Not significant (n.s.), \*\*\**p* < .001 and \*\*\*\**p* < .0001.

Because the proteomic clusters were associated with distinct histological and microenvironmental states, we next asked whether the RNA-defined groups also differed in their inferred cellular composition. By performing cell enrichment analysis, we observed enrichment of osteoblast and fibroblast signatures in Clusters B and C (**Figure 6E)**. We examined the expression of KLK3 and TPD52, together with KRT8 and KRT18, epithelial keratins highly expressed by PC cells, but not by the normal bone microenvironment. Both keratin genes were expressed at significantly lower levels in RNA-defined Cluster A than in Clusters B and C (**Figure 6F**), suggesting that Cluster A contains relatively low tumor content and a greater contribution from preserved bone marrow and immune cell populations. By mapping our proteomic signatures onto independent transcriptomic datasets, we demonstrated that the molecular and cellular features identified in our cohort were also evident at the RNA level. The transcriptomic groups showed molecular states that closely paralleled those of the proteomic clusters, supporting a shared continuum of PCBM phenotypes.

### PCBM markers defining sclerotic lesions discriminate bone metastases from soft tissue metastases in transcriptomic data

To further refine the list of 58 differentially expressed proteins that segregates the PCBMs, we used a random forest approach to identify the 8 proteins that contributed the most to the segregation of the osteosclerotic subtypes (**Figure 7A, Supplemental Figure 7**). In order to evaluate whether this ECM-representative protein panel has translational potential, we assessed whether it could distinguish bone metastases from soft-tissue metastases in the three independent mCRPC RNA-seq datasets: WCDT, PROMOTE, and the ECDT (**Figure 7B**). We first performed PCA using all detected genes across all metastatic samples (**Figure 7C**). This unsupervised analysis showed little separation of lesions based on metastatic sites. In contrast, PCA restricted to the 8 osteosclerotic markers clearly separated bone metastases, and a subset of adrenal metastases from soft-tissue metastases (**Figure 7D**), demonstrating that this marker panel captures transcriptional features specific to the PCBM microenvironment.

**Figure 7.**
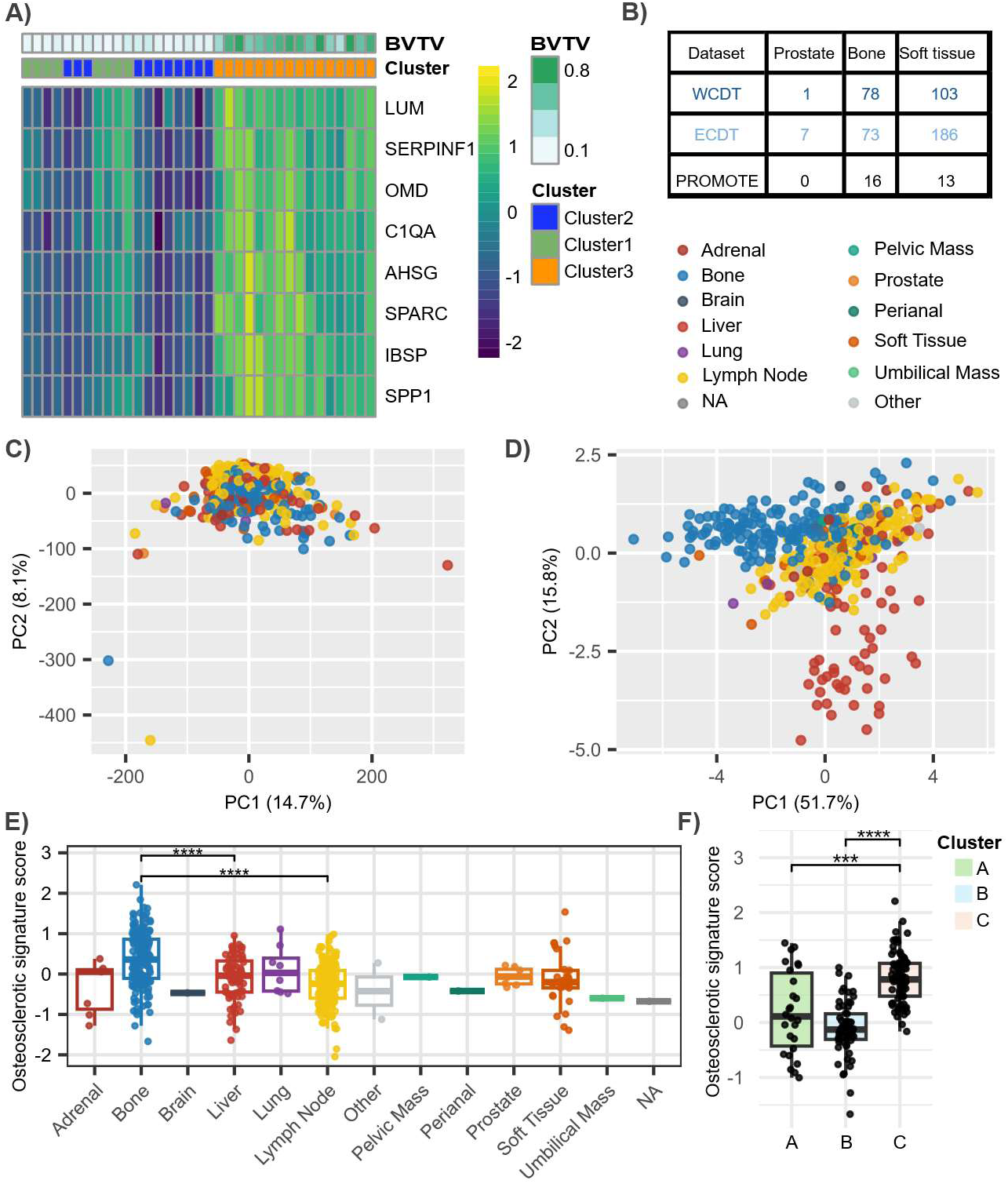
Validation of biomarker panel in independent RNA-seq datasets demonstrates high predictive power. **(A)** Expression of the 8 osteosclerotic marker proteins in proteomic data B) External validation cohorts used with transcriptomic data for prostate tumours, bone metastasis, and soft tissue metastasis. (C) PCA of combined RNA-seq data (ECDT, WCDT and PROMOTE) of all genes. (D) PCA of the combined dataset restricted to the osteosclerotic marker panel. (E) Osteosclerotic signature score across metastasis sites. Significance level only shown for comparisons between bone with liver and lymph node. (F) Distribution of osteosclerotic scores by RNA clusters indicated by color. \*\*\**p* < .001 and \*\*\*\**p* < .0001.

We next calculated an osteosclerotic signature score for each sample based on the 8 markers. Bone metastases exhibited significantly higher osteosclerotic signature scores than metastases from all other anatomical sites (**Figure 7E**). Furthermore, when bone metastases were stratified according to the transcriptional clusters defined in Figure 4, samples belonging to Cluster C with enriched extracellular matrix organization and bone remodeling pathways displayed significantly higher osteosclerotic signature scores than the other cluster (**Figure 7F**). Our findings demonstrate that unique molecular features of PCBMs are conserved across proteomic and transcriptomic datasets and can robustly segregate PCBMs from other metastatic lesion types, and that a select group of PCBM molecular features might be translatable as a signature of osteosclerotic PCBM.

## Discussion

The devastating consequences of PCBM on patients’ quality of life are the consequence of structural and molecular changes in bone. PCBM are heterogeneous skeletal lesions that are characterized by simultaneous osteoblastic and osteolytic remodeling. Although these lesions are radiographically classified as predominantly osteosclerotic, they exhibit irregular bone deposition, increased porosity, disorganized collagen architecture, and reduced mechanical integrity, features that resemble woven bone generated during skeletal development and fracture repair (6). Furthermore, interactions between PC and bone are critical in PC progression, but clinically applicable aspects of these mechanisms remain to be developed (23–25). In this study, we define molecularly distinct PCBM subtypes that closely associate with distinct histological and structural states from relatively preserved marrow to tumor-infiltrated and extensively osteosclerotic lesions. We also established a marker signature of osteosclerotic remodeling that is reproducible across independent metastatic PC datasets and reflects progressive changes within the bone microenvironment. These findings reflect tumor-driven remodeling of the bone microenvironment and establish molecular classification that may inform targeted therapeutic strategies and biomarker development.

The spectrum of proteomic states described here aligns with the functional declines required for the establishment and survival of PC in the bone environment. Neutrophils account for approximately 60% of resident bone marrow immune cells (26), and they play an important role in the early disease progression of PCBM (27, 28). Bone marrow neutrophils isolated early in disease were directly cytotoxic to PC cells, and experimental neutrophil depletion accelerated tumor growth within bone. More broadly, as PC cells establish in the bone environment, they alter the immune structure of the local and peripheral sites, creating a more immunologically “cold” niche (27, 29). This progression mirrors our proteomic data, in which the loss of the neutrophil-marked signature from Cluster 1 through Cluster 2 and Cluster 3 parallels the transition from a tumor-restrictive to an immunosuppressive marrow environment. Our observations support a model in which PCBM develops through progressive disruption of the marrow immune and stromal environment, followed by tumor establishment and extensive osteosclerotic remodeling. Nevertheless, the cross-sectional nature of the present cohort does not establish a temporal sequence between these lesion states. Representative temporal modeling will be required to determine whether these proteomic states represent sequential stages of disease progression and whether they are generalizable across patients.

We observed extensive COL III deposition within the osteosclerotic lesions. COL III-positive fibres extended throughout the marrow space, accumulated along bone surfaces, and surrounded or partially encapsulated tumor cell clusters. This pattern is consistent with the development of a fibrotic stromal response within the PCBM microenvironment. Excessive COL III deposition is a well-established feature of primary myelofibrosis, a myeloproliferative neoplasm characterized by progressive bone marrow fibrosis, neo-angiogenesis, disruption of normal haematopoiesis, and, in advanced disease, osteosclerosis (30). In primary myelofibrosis, bone marrow fibrosis follows a defined structural progression: COL III initially forms a reticular fiber network within the marrow, which is later reinforced by thicker, more densely packed COL I fibers as fibrosis advances toward overt osteomyelofibrosis and osteosclerosis (31). In its most severe form, this fibrotic cascade culminates in extensive sclerotic bone formation, offering a histopathological parallel to the sclerotic bone formation seen in PCBM (32, 33). Although the initiating mechanisms differ between primary myelofibrosis and PCBM, our observations of an increased COL III matrix forming excessive reticular fibres which occupy most of the bone marrow spaces, and subsequent COL I deposition and mineralization, suggests the presence of common mechanisms of bone marrow fibrosis, which may represent interconnected components of a shared, maladaptive tissue-remodelling response. Importantly our observations highlight the need to understand PCBM as a multidimensional disease state with profound effects on skeletal health, bone marrow homeostasis, and tumor burden. Deciphering the cellular and molecular mechanisms of this process are of paramount relevance for the comprehensive understanding of metastatic prostate cancer biology, and is a critical need to develop therapeutic strategies and pain relief for PC patients.

Cancer-associated fibroblasts (CAFs) are critical in cancer pathogenesis with their ability to develop de novo steroidogenesis, enhance neovascularization, elicit invasiveness, and suppress anti-tumor immunity to promote tumorigenesis, and support an immunosuppressive environment (34–39). Single-cell RNA-seq data from PCBM samples have identified osteoblast subtypes molecularly resembling CAFs, and a distinct population of EMP1+/COL3A1+ fibroblasts has been shown to be enriched in bone metastases from prostate, breast, and renal cancer, where fibroblast-derived COL3A1 signals to tumor cells via the ADGRG1 receptor to potentially facilitate the metastatic process (40, 41). Consistent with these observations, our transcriptomic analysis demonstrated co-enrichment of fibroblast and tumor-associated signatures. Taken together with the increased fibrosis and COL III deposition observed in our PCBM specimens, we speculate that CAFs, or CAF-like fibroblasts contribute to the fibrotic remodeling associated with osteosclerosis in PCBM, potentially representing an early step in the development of aberrant osteosclerotic bone in PCBM.

In addition to COL III, our proteomic analysis identified increased expression of several bone-remodelling proteins in osteosclerotic lesions, including the IBSP and SPP1. Both are well-characterized regulators of osteoblast differentiation, mineralization, and bone turnover, and have also been implicated in tumor–bone interactions (20). IBSP and SPP1 expression are increased in prostate and breast cancer bone metastases, while exposure to IBSP or SPP1 can enhance tumor-cell adhesion, migration, and invasion (42, 43). Consistent with these observations, our IHC demonstrated IBSP and SPP1 expression in tumor cells as well as osteoblasts and osteocytes associated with pathological bone. IBSP is expressed during early bone deposition and promotes mineral formation by binding collagen, calcium, and hydroxyapatite at sites of active bone formation with its enrichment in rapidly deposited woven bone further supporting its association with active pathological remodelling (44, 45). SPP1 is similarly enriched at sites of active mineralization but primarily regulates crystal growth and also promotes osteoclast attachment to mineralized matrix, linking it to bone resorption and turnover (46, 47). The convergent expression of IBSP or SPP1 in the PCBM niche therefore suggests a potential link between tumor–bone crosstalk and an exaggerated bone-healing response that promotes disorganized osteosclerotic matrix formation.

From a disease outcomes perspective, we did not identify a significant association between the molecular clusters and overall patient survival (**Supplementary Figure 8A**). This may be due to the substantial spatial heterogeneity of PCBM since each analyzed specimen represents only a localized region of a patient’s overall disease burden (48). Consistent with this, specimens obtained from different skeletal sites within the same patient can present as different PCBM subtypes, and here, were assigned to different proteomic clusters (**Supplemental Figure 8B**) demonstrating that distinct disease states can coexist within an individual (6). Consequently, the molecular phenotype of a single bone lesion may not accurately reflect the systemic extent or severity of disease and may therefore have limited prognostic value for patient-level outcomes. Accurate prediction of clinical outcomes will likely require approach that captures systemic changes associated with the disease severity. Our mass spectrometry approach was also strongly influenced by the abundance of dense extracellular matrix within PCBM lesions, which limited our ability to resolve tumor cell-specific protein expression and define associations between PCBM lesion types and tumor types. This is a meaningful limitation, since PC develops and evolves in close conjunction with its surrounding stroma, and reciprocal signaling between tumor and stromal compartments is known to promote malignant progression, survival, and therapy resistance (49). Future work incorporating tumor-cell–resolved proteomic or spatial approaches will therefore be important to understand the heterogeneity of tumor cell populations across PCBM lesion types, and how differences in tumor molecular phenotypes shape the resulting bone lesion type.

## Methods

### Sex as a biological variable

Patient samples included in this study were exclusively from male patients, as prostate cancer is a male-specific disease.

### Study approval

All described procedures were approved by the institutional review boards at University of Washington (2341) and University of British Columbia (H21-02668). Cadaveric thoracic and lumbar vertebral cores and control samples were obtained as previously described (6). Samples were processed for histological analysis as previously described (50).

### Proteomics Analysis

Proteins were extracted from formalin-fixed paraffin-embedded demineralized bone sections (5 x 5 µm sections). TMT labeling (10plex) was carried out following manufacturers’ instructions (ThermoFischer Scientific). Peptide detection was performed using an Orbitrap Fusion Lumos MS platform (Thermo Scientific). Data were acquired using a data-dependent acquisition method and processed in Proteome Discoverer 2.5.0.400 using Sequest HT against a human SwissProt database (TaxID 9606; January 2022 release). Protein abundances were normalized to a pooled TMT reference channel, and statistical significance was assessed using the software’s background-based ANOVA test.

### Differentially expression analysis

Proteomic data analysis was conducted in R (v4.4.1). Proteins with more than 50% missing values were excluded from downstream analyses. Missing values for the remaining proteins were imputed using the k-nearest neighbors method. Protein intensity values were log-transformed and subsequently median normalized. PCA was performed using the stats package (version 4.4.1). Heatmaps with hierarchical clustering were created using the pheatmap package (version 1.0.13). Differential expression analysis was conducted using a linear model from the limma package (version 3.62.1). P-values were adjusted for multiple comparisons using the Benjamini-Hochberg method. Proteins with a fold change > 1.5 and an adjusted p-value < 0.05 were considered statistically significant.

### Immunohistochemistry

The antibodies used in this study are listed in Supplemental Table 2. Antigen retrieval was performed on mounted tissue sections in 10 mM citrate buffer (pH 6.0) overnight in a 60 °C water bath. Endogenous peroxidase activity was quenched using 3% H₂O₂ for 30 min, followed by blocking of non-specific protein binding with 3% bovine serum albumin in PBS for 60 min. Sections were then incubated with primary antibody at 4°C overnight. After washing with PBS, sections were incubated with the appropriate horseradish peroxidase-conjugated reagent (**Supplemental Table 2**) for 1 h at room temperature. Immunoreactivity was visualized using an Abcam 3,3′-diaminobenzidine (DAB) chromogenic detection kit according to the manufacturer’s instructions.

### Functional Analysis

Significant proteins identified in the differential expression analysis were analyzed for over-representation in Gene Ontology biological processes and Reactome pathways using the gprofiler2 package (51). The g:SCS method, built into gprofiler2, was used to correct for multiple testing. A corrected p-value < 0.05 was considered statistically significant.

### Batch-effect correction

Publicly available RNA sequencing datasets from three mCRPC patient cohorts, including WCDT, ECDT, and PROMOTE cohorts, were combined for analysis. Batch effects were corrected with ComBat following cohort integration to minimize technical variation between cohorts (52).

### RNA data analysis

The WCDT and the PROMOTE mCRPC transcriptomic datasets were obtained from the Quigley Lab public data repository, cBioPortal, and the Gene Expression Omnibus. Processed RNA-seq data from WCDT and PROMOTE were downloaded from the Quigley Lab repository as transcripts per kilobase million (TPM) values (53, 54). ECDT mCRPC cohort was accessed through cBioPortal; fragments per kilobase of transcript per million mapped reads values were converted to TPM prior to analysis (55). Expression matrices were then harmonized by gene symbol, log-transformed, and annotated by metastatic tissue site where available. GSEA was performed in R using gene sets obtained from the Molecular Signatures Database (MSigDB), with Hallmark pathways used for functional pathway enrichment analysis. Cell enrichment analysis was done using xCell (56).

## Supporting information

Supplemental data files, for linking supporting figures and tables on the preprint site

Supporting data vales, will be used to link proteomic data file on the prepint site

## Data availability

Data supporting the finding of the study is available in supplemental tables. No original code was created in this study.

## Statistical analysis

For comparisons across multiple groups, statistical significance was assessed using the Kruskal–Wallis test. Where applicable, pairwise group comparisons were subsequently performed using Wilcoxon rank-sum tests with adjustment for multiple comparisons. Statistical analyses and visualization were performed in R.

## Author contributions

Conceptualization, DX, CM, EC, LDT, RW, FE, and MEC; Research design, DX, FE, and MEC; Bioinformatic analysis, DX and CC; Histological staining, DX, KC, and BM; Sample processing, BM, DL, and NJ; Immunohistochemistry, KC; Figure generation, DX, KC; Proteomics experiments, HA; Provision of samples, CM, EC, and LDT; Writing original draft, DX and FE; Writing – review and editing, all authors.

## AI disclosure

ChatGPT (OpenAI; GPT-5.6 Sol) was used between September and October 2026 to assist with manuscript for language editing and refinement of manuscript text. The authors assume full responsibility for ensuring the integrity and accuracy of AI output presented in this article.

## Acknowledgements

We thank the patients and their families, Pete Nelson, Evan Yu, Elahe Mostaghel, Heather Cheng, Celestia Higano, Bruce Montgomery, Mike Schweizer, Andrew Hsieh, Jonathan Wright, Daniel Lin, Funda Vakar-Lopez, Xiaotun Zhang, Martine Roudier, Paul Lange, Robert Vessella,, and the rapid autopsy teams for their contributions to the University of Washington Prostate Cancer Donor Rapid Autopsy Program. We thank the donors and families of the University of British Columbia Body Donor Program, and the contributions of Matthew Tinney, Grant Regier, Manouchehr Madani Civi, Adel Hajjay, Lexi Busse, Edwin Moore. We thank Danmei Liu and Doris Liang for their assistance with sample processing. We are grateful to patients with lived experiences, Mark White, Robert Wong, John Stewart, Len Gross for their guidance throughout this work. The authors acknowledge that this work was performed in the unceded traditional territories of the xʷməθkʷəy̓əm (Musqueam), Sḵwx̱wú7mesh (Squamish), and səlilwətaɬ (Tsleil-Waututh) Nations.

