## Supplemental data files, for linking supporting figures and tables on the preprint site for "Proteomic Characterization of Prostate Cancer Bone Metastasis Subtypes Defines Progressive Remodeling States of the Bone-Tumor Microenvironment"

**Supplemental Table 1. Bone volume fraction across proteomic clusters**

| Cluster | N | BV/TV, mean ± SD | Median (Q1–Q3) | Range |
|---|---|---|---|---|
| Cluster 1 | 8 | 0.109 ± 0.035 | 0.101 (0.091–0.116) | 0.071–0.182 |
| Cluster 2 | 11 | 0.130 ± 0.037 | 0.126 (0.103–0.153) | 0.082–0.204 |
| Cluster 3 | 16 | 0.608 ± 0.148 | 0.639 (0.529–0.705) | 0.321–0.801 |

**Supplemental Table 2. Antibody used for immunohistochemistry**

| Antibody | Cat # | Species | Brand | Dilution |
|---|---|---|---|---|
| IBSP | A4222.2 | Rabbit | Immundiagnostik AG | 1:2000 |
| Collagen III | AB6310 | Mouse | Abcam | 1:2000 |
| Collagen 1 | BS-0578R | Rabbit | Bioss | 1:2000 |
| SPP1 | A4226AG.1 | Synthetic | Immundiagnostik AG | 1:2000 |
| Mouse Igg | 205719 | Goat | Abcam | 1:1000 |
| Rabbit Igg | 205718 | Goat | Abcam | 1:1000 |

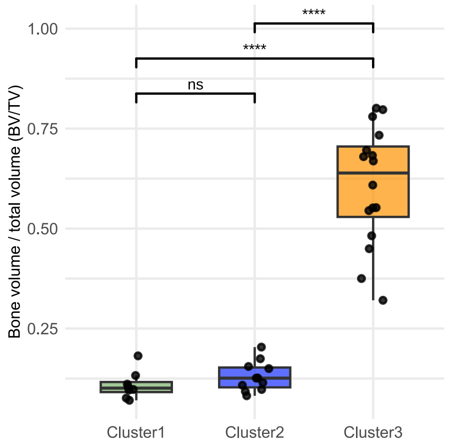


**Supplemental Figure 1. Bone volume fraction across proteomic clusters.** Boxplots show the median and interquartile range, with individual samples overlaid as points. Pairwise differences between clusters were assessed using Wilcoxon rank-sum tests with Benjamini–Hochberg correction for multiple comparisons. Significance is indicated as ns, not significant; **** $P < 0.0001$.

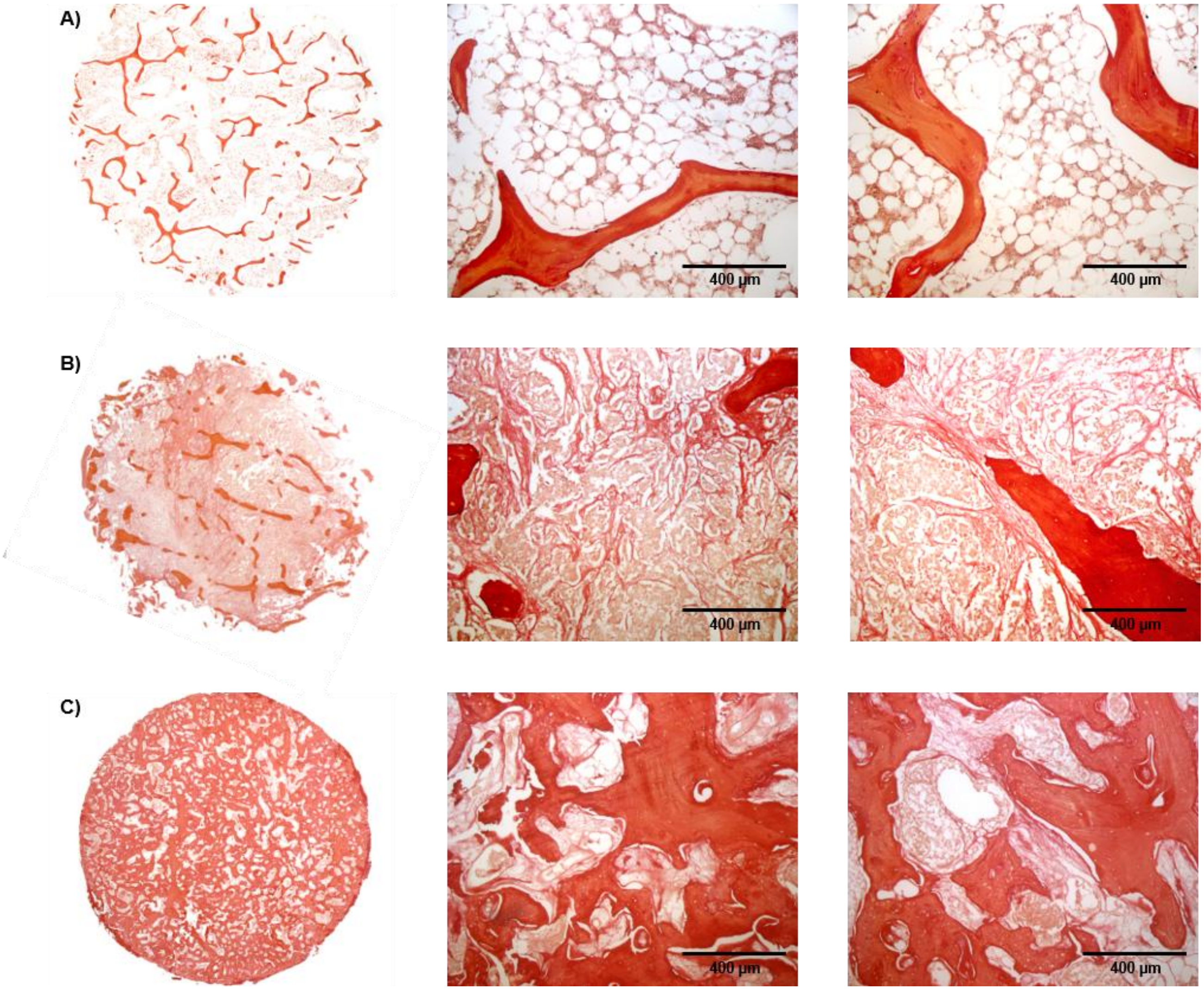


**Supplemental Figure 2. Sirus Red staining from samples across proteomic clusters.** (A-C) Representative Sirius Red-stained sections from Cluster 1 (A), Cluster 2 (B), and Cluster 3 (C).

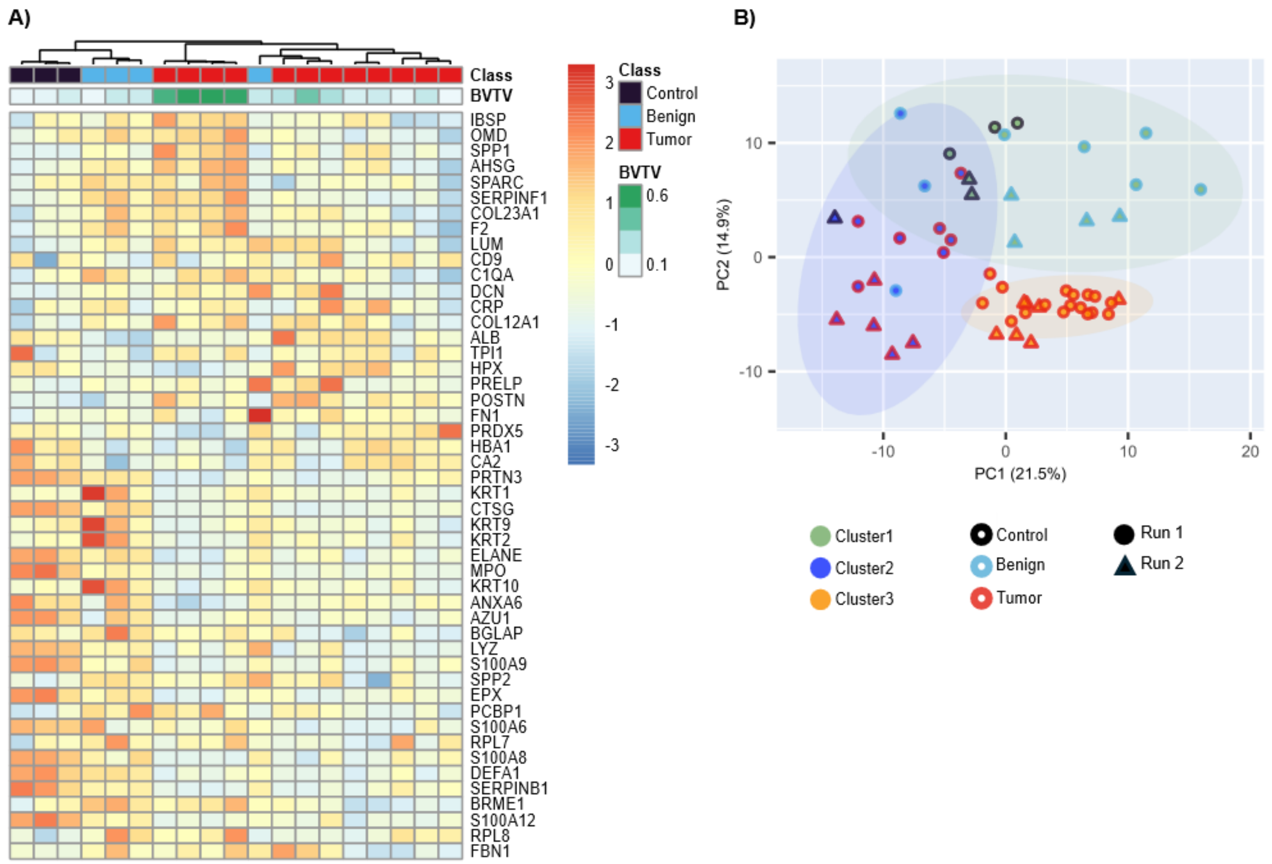


**Supplemental Figure 3. Validation of proteomic cluster signatures in an independent sample set.** (A) Relative protein expression of proteins uniquely upregulated in each proteomic cluster in an additional set of 19 PCBM samples. Annotation bars indicate BV/TV and sample type. (B) Combined PCA of the initial 35 samples (Run 1) and the additional 19 samples (Run 2), based on the cluster-associated protein marker panel. Point fill indicates proteomic cluster, point outline indicates sample type, and point shape indicates MS run.

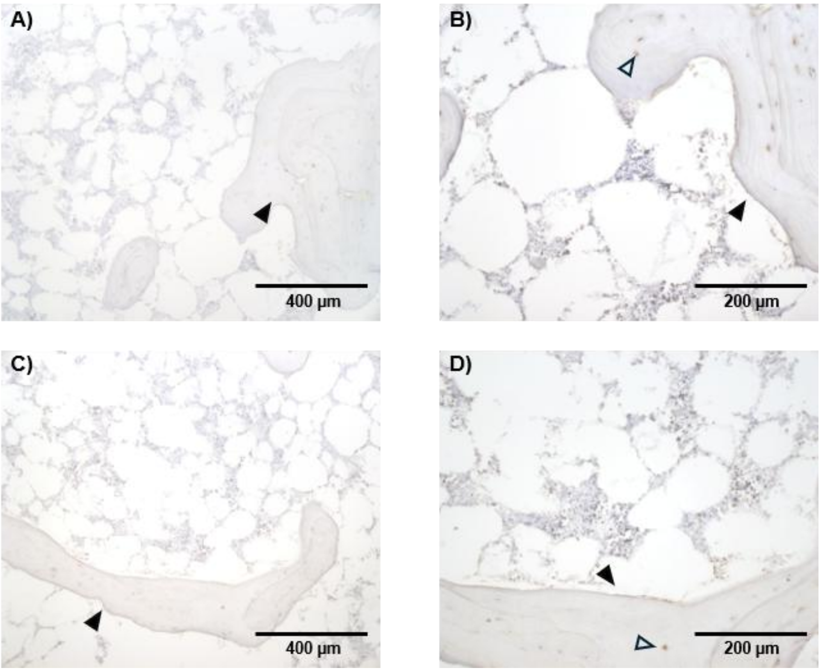


**Supplemental Figure 4. Bone sialoprotein staining in control bone marrow.** (A, C) Representative images of control bone marrow at 10× magnification. (B, D) Corresponding higher-magnification views (20×) of the regions shown in A and C. Arrowheads indicate trabecular bone, and open arrowheads indicate osteocytes.

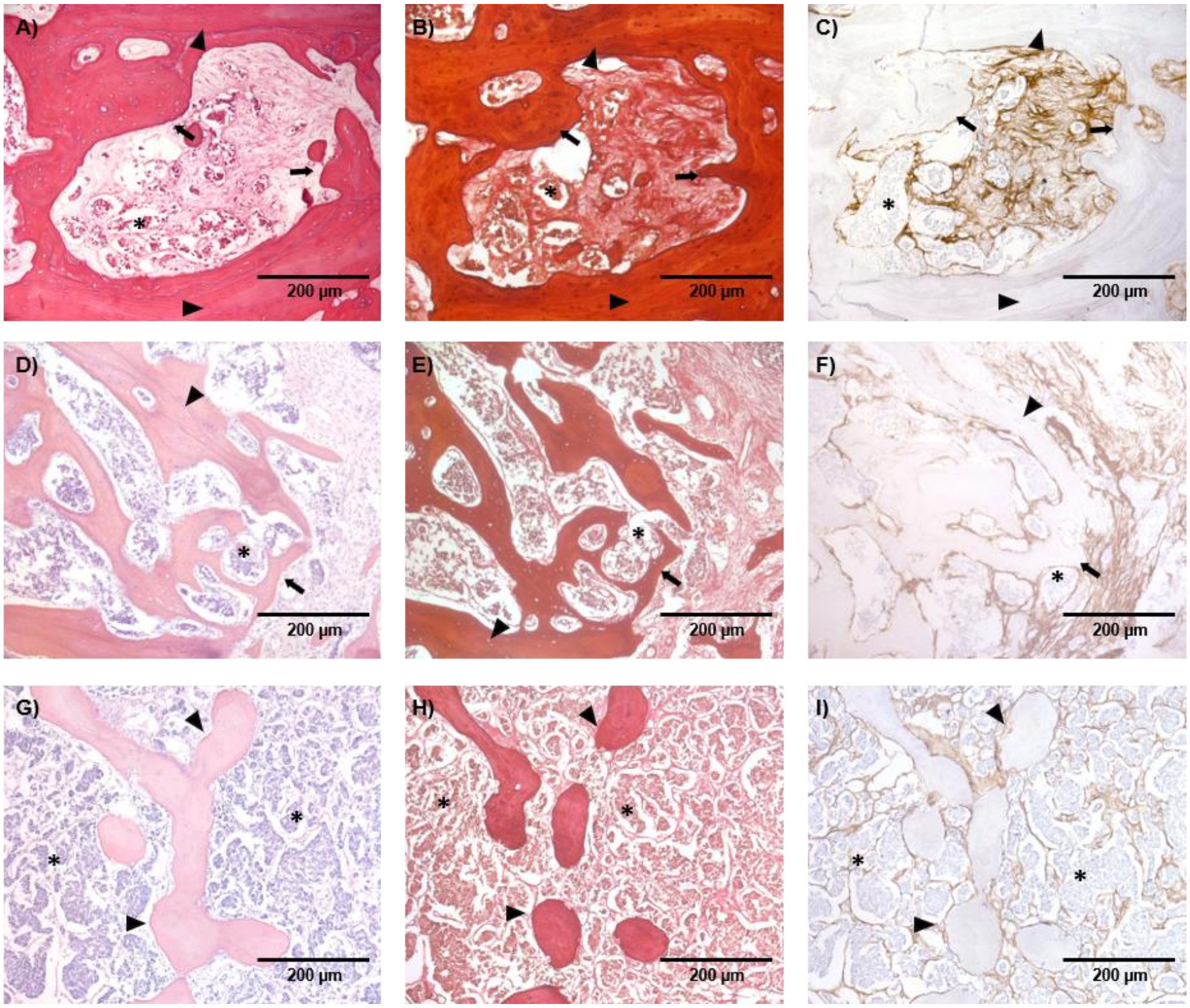


**Supplemental Figure 5. Collagen III deposition patterns in PCBM.** (A, D, G) Representative H&E-stained sections showing distinct patterns of collagen III deposition in PCBM at 20× magnification. (B, E, H) Corresponding Sirius Red staining. (C, F, I) Corresponding COL III immunohistochemical staining. Residual trabeculae bone (arrowheads), osteosclerotic bone (arrows), and tumor cells (asterisks).

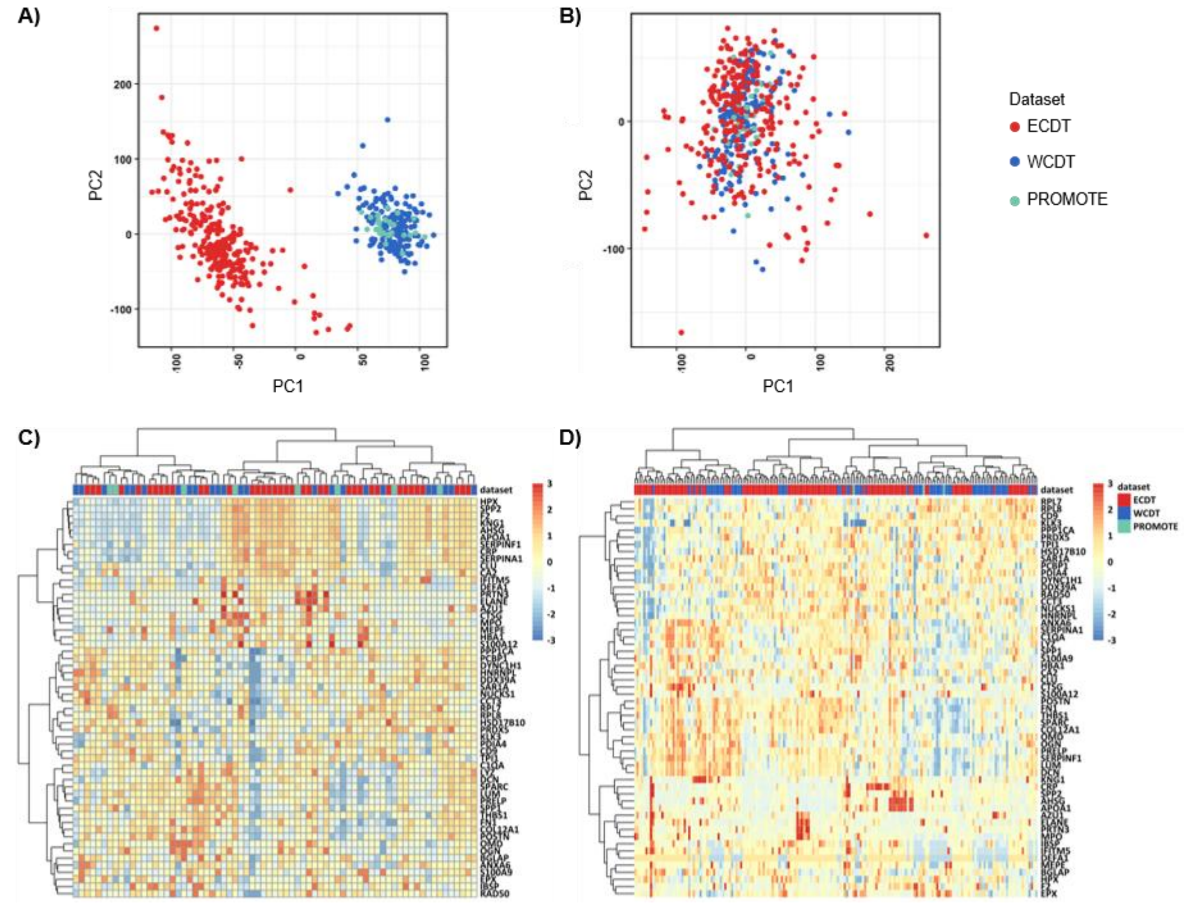


**Supplemental Figure 6. Batch correction and transcriptomic clustering of metastatic prostate cancer RNA-seq cohorts.** (A) PCA of RNA-seq data from the ECDT, WCDT, and PROMOTE cohorts before batch correction. (B) PCA of the merged dataset after batch correction. (C) Unsupervised hierarchical clustering heatmap of proteomics-derived marker genes in adrenal metastasis samples from the merged RNA-seq dataset. (D) Unsupervised hierarchical clustering heatmap of proteomics-derived marker genes in lymph node metastasis samples from the merged RNA-seq dataset.

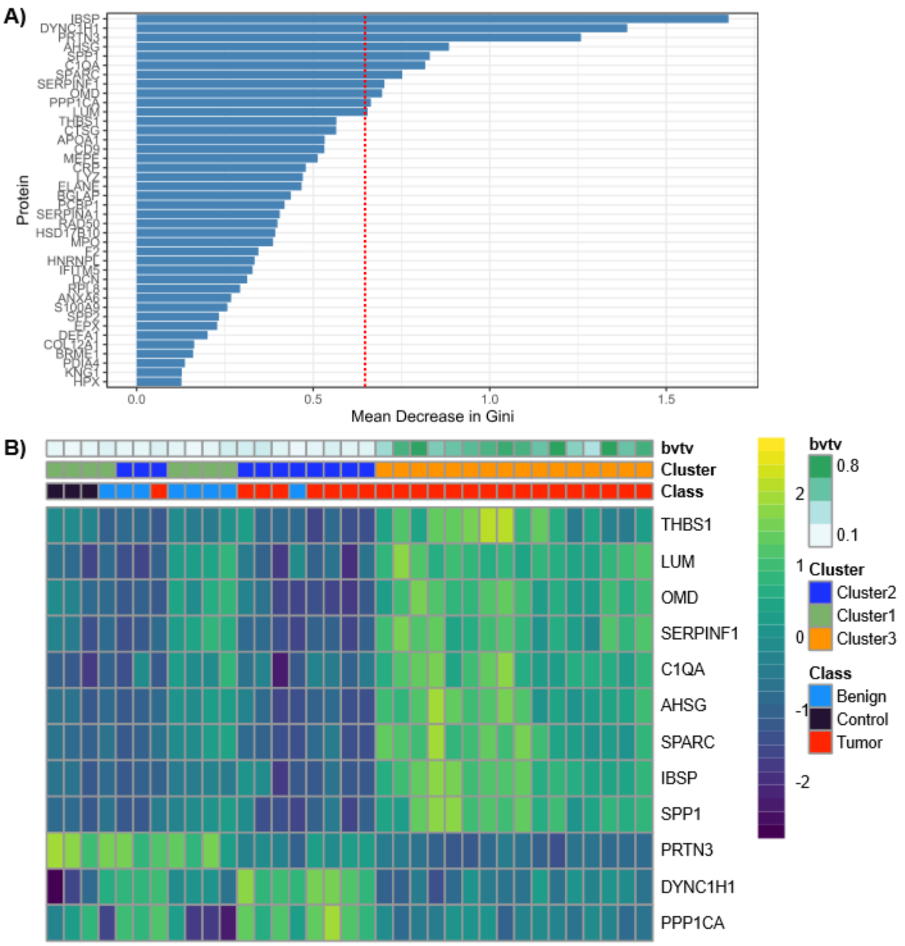


**Supplemental Figure 7. Random forest-derived marker selection across proteomic clusters.** (A) Random forest variable-importance plot showing the top proteins distinguishing Cluster 3 from Clusters 1 and 2, ranked by mean decrease in Gini index. The red dotted line indicates the threshold used for marker selection. (B) Heatmap of the selected random forest marker proteins across samples.

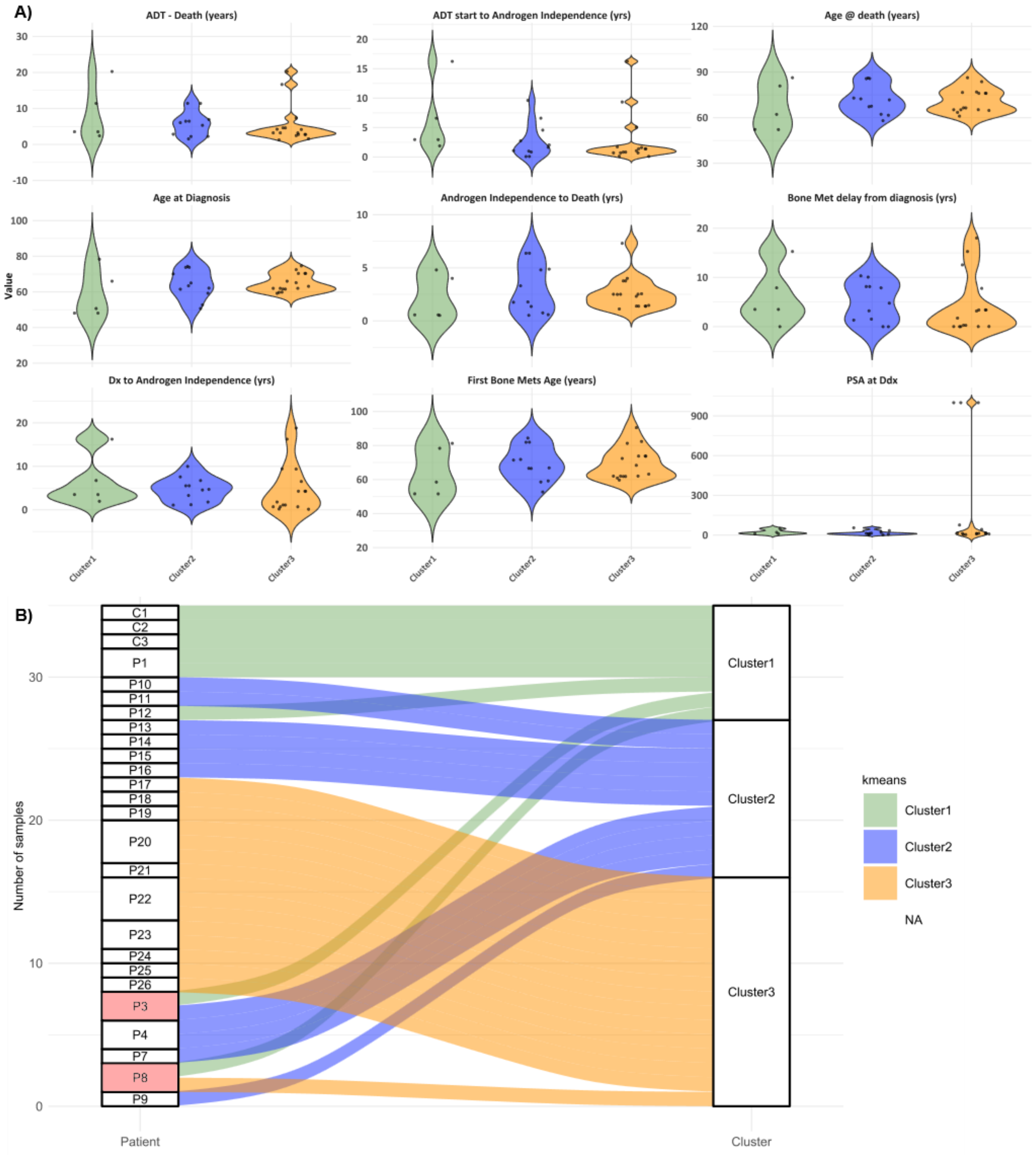


**Supplemental Figure 8. Patient distribution and clinical features across proteomic clusters.** (A) Violin plots showing the distribution of selected clinical features across the three proteomic clusters, with individual samples overlaid as points. (B) Alluvial plot showing the distribution of patient samples across the three proteomic clusters. Patients with samples assigned to more than one cluster are highlighted in pink.
